# Integration of immunome with disease-gene network reveals common cellular mechanisms between IMIDs and drug repurposing strategies

**DOI:** 10.1101/2019.12.12.874321

**Authors:** Abhinandan Devaprasad, Timothy RDJ Radstake, Aridaman Pandit

## Abstract

**Objective:** Development and progression of immune-mediated inflammatory diseases (IMIDs) involve intricate dysregulation of the disease associated genes (DAGs) and their expressing immune cells. Due to the complex molecular mechanism, identifying the top disease associated cells (DACs) in IMIDs has been challenging. Here, we aim to identify the top DACs and DAGs to help understand the cellular mechanism involved in IMIDs and further explore therapeutic strategies.

**Method:** Using transcriptome profiles of 40 different immune cells, unsupervised machine learning, and disease-gene networks, we constructed the Disease-gene IMmune cell Expression (DIME) network, and identified top DACs and DAGs of 12 phenotypically different IMIDs. We compared the DIME networks of IMIDs to identify common pathways between them. We used the common pathways and publicly available drug-gene network to identify promising drug repurposing targets.

**Result:** We found CD4^+^Treg, CD4^+^Th1, and NK cells as top DACs in the inflammatory arthritis such as ankylosing spondylitis (AS), psoriatic arthritis, and rheumatoid arthritis (RA); neutrophils, granulocytes and BDCA1^+^CD14^+^ cells in systemic lupus erythematosus and systemic scleroderma; ILC2, CD4^+^Th1, CD4^+^Treg, and NK cells in the inflammatory bowel diseases (IBDs). We identified lymphoid cells (CD4^+^Th1, CD4^+^Treg, and NK) and their associated pathways to be important in HLA-B27 type diseases (psoriasis, AS, and IBDs) and in primary-joint-inflammation-based inflammatory arthritis (AS and RA). Based on the common cellular mechanisms, we identified lifitegrast as potential drug repurposing candidate for Crohn’s disease, and other IMIDs.

**Conclusion:** Our method identified top DACs, DAGs, common pathways, and proposed potential drug repurposing targets between IMIDs. To extend our method to other diseases, we built the DIME tool. Thus paving way for future (pre-)clinical research.

## 1. Introduction

The genetic and epigenetic heterogeneity has been known to play a major role in the development and progression of complex diseases. The past two decades has seen a major surge in studies that characterize genes and loci associated with diseases [1]. The use of high-throughput omics technology and functional screenings have boosted our knowledge about genetic, epigenetic and metabolic factors underlying complex diseases [1]. As a result of these genetic and epigenetic screenings, we now know that the majority of complex diseases and genes/loci have a many-to-many relationship meaning that a complex disease is linked to many different genes and a gene/loci might be associated with many different diseases [2]. Large high-throughput screening studies have typically used bulk tissue or whole blood to study disease associated genes (DAGs). However, the expression of each gene is known to vary between tissues and cell types [3,4]. Thus, bulk tissue- or blood-based studies on DAGs do not consider the role played by different cells and tissues in the disease biology. To improve the understanding and molecular basis of complex diseases, a large number of research groups and consortiums have started to functionally identify disease associated cells (DACs) or tissue types [3–7]. The Genotype-Tissue Expression (GTEx) is one such valuable project, which maps gene expression profiles of 54 different human tissue types and the corresponding expression quantitative trait loci (eQTLs) [5–7]. Furthermore, the growth of single cell technologies have advanced our understanding of DACs and have helped in identifying cell types associated with complex diseases including cancer [8], Alzheimer’s [9], rheumatoid arthritis [10], among others.

The immune system is known to play a key role in the development and progression of immune-mediated as well as non-immune mediated chronic diseases. A large number of association and functional studies have shown that multiple DAGs are expressed in immune cells and perturbing these DAGs can modulate immune cell functions [11]. However, very few studies have explored the impact of DAGs on specific cell types and even fewer on immune cells, many of which focus on limited number of cell subsets [12–16]. Recently Schmiedel *et al*. studied the effect of genetic variants on the expression of genes in 13 different immune cell types [17]. However, this study largely focused on the analysis of genetic variants and their impact on a total of 13 immune cell types: monocytes (classical and non-classical), NK cells, naïve B-cells and nine sub-populations of T-cells. Immune-mediated inflammatory diseases (IMIDs) are complex in nature, with the involvement of several different types of immune cells. For example, in rheumatoid arthritis, the immune cells such as B-cells, T-cells, macrophages, mast cells, dendritic cells, and NK cells are known to play a major role in the pathogenesis of the disease [18]. Insights on the exact mechanism of action is crucial for developing successful therapies for the disease. This becomes particularly challenging for IMIDs due to the involvement of several cell types. The massive undertaking of GWAS for the IMIDs have enabled mapping of some of the molecular mechanisms of the IMIDs [19–22]. However, most of these have uncovered only the tip of the iceberg and further research is required to understand the etiology of these diseases with respect to the several different immune cells at play, and to identify any mechanistic overlap between the IMIDs. This approach of identifying the key immune cells at play and their mechanism in the IMIDs would set a robust rationale for exploring therapeutic strategies.

In this study, we mapped the largest available and expert curated disease-gene network (from the DisGeNet curated from 16 different databases) [23] on the largest *immunome* data comprising gene expression profiles of 40 different immune cell types, curated by us. We further built a tool using an unsupervised machine learning algorithm, the disease-gene network, and the *immunome* to create the Disease-gene IMmune cell Expression (DIME) network. Hereby, the tool is referred to as the DIME; the analysis using this tool is referred to as the DIME analysis. Using DIME, we then quantified the effects of 3957 DAGs on the *immunome*, to identify DACs for 12 phenotypically different IMIDs. We used the DIME to: (1) study the underlying cell-specific mechanisms; (2) identify common DACs and their top weighted DAGs (hereby referred to as common cell-gene network) between different pairs of diseases; and (3) identify drug repurposing targets using the common cell-gene network. The DIME is available as a user-friendly R tool (https://bitbucket.org/systemsimmunology/dime), to identify the top genes and cells associated with the disease of interest for: (1) diseases from the DisGeNet, (2) diseases from the EBI genome wide association study (GWAS) catalogue, or (3) custom set of genes defined by the user.

## 2. Methods

### 2.1. Transcriptome data - *Immunome*

The transcriptome data consists of RNA-sequencing datasets of 40 different immune cell types curated using 316 samples from a total of 27 publicly available datasets (see Supplementary Table 1 for list of GEO datasets and samples used). The 40 different immune cells cover the entire hematopoietic stem cell differentiation tree comprising of 9 progenitors, 19 lymphoid, and 12 myeloid cell types. The samples used here were manually curated considering only the unstimulated (except for macrophages, that were monocyte derived) immune cells that were sorted using Fluorescence-activated cell sorting (FACS) and were isolated from either blood, bone marrow or cord blood from healthy donors. The processed, batch corrected, and normalized data of the 40 immune cells is referred here as the *immunome* (see Supplementary methods for details).

**Table 1:**
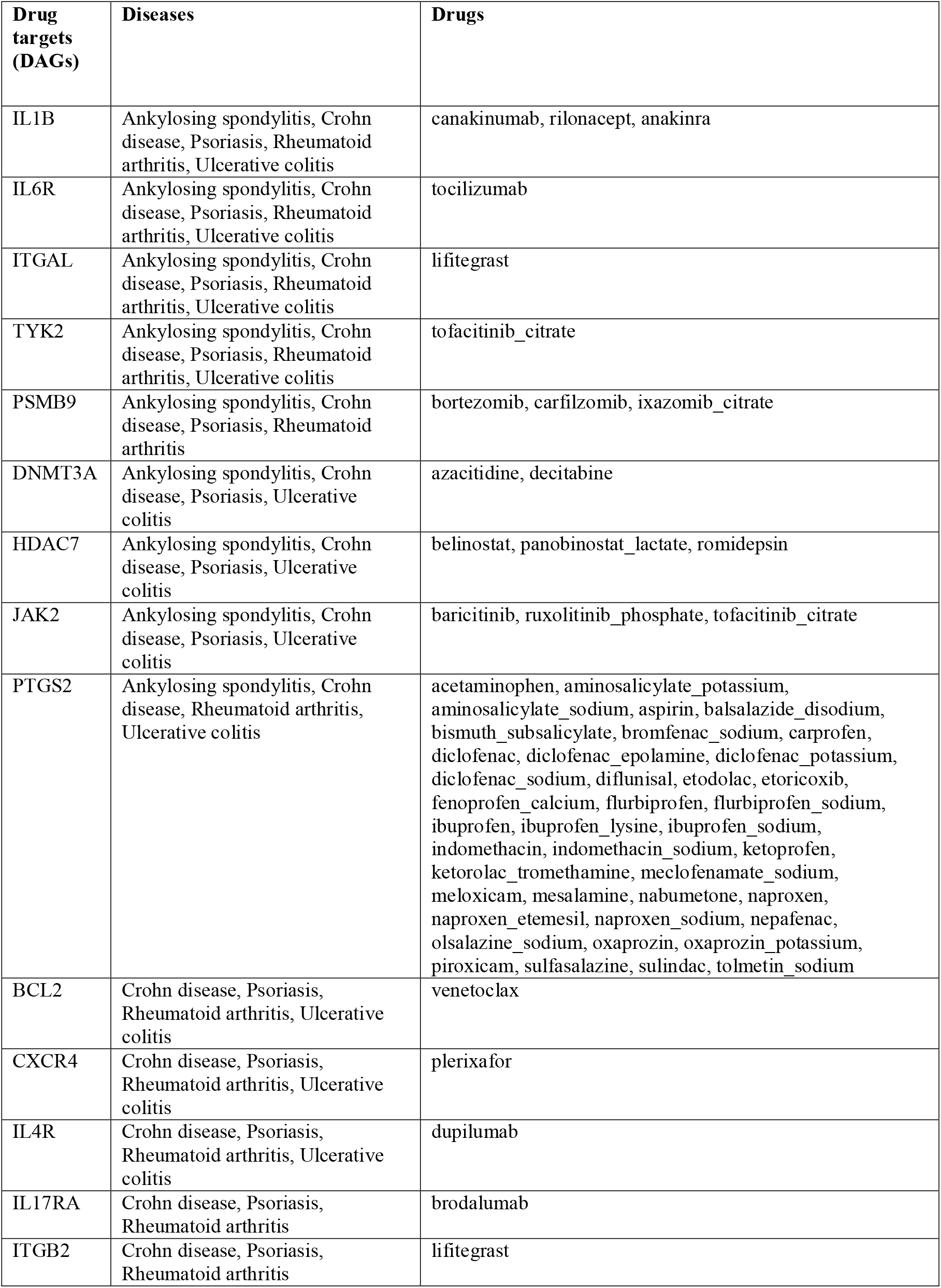
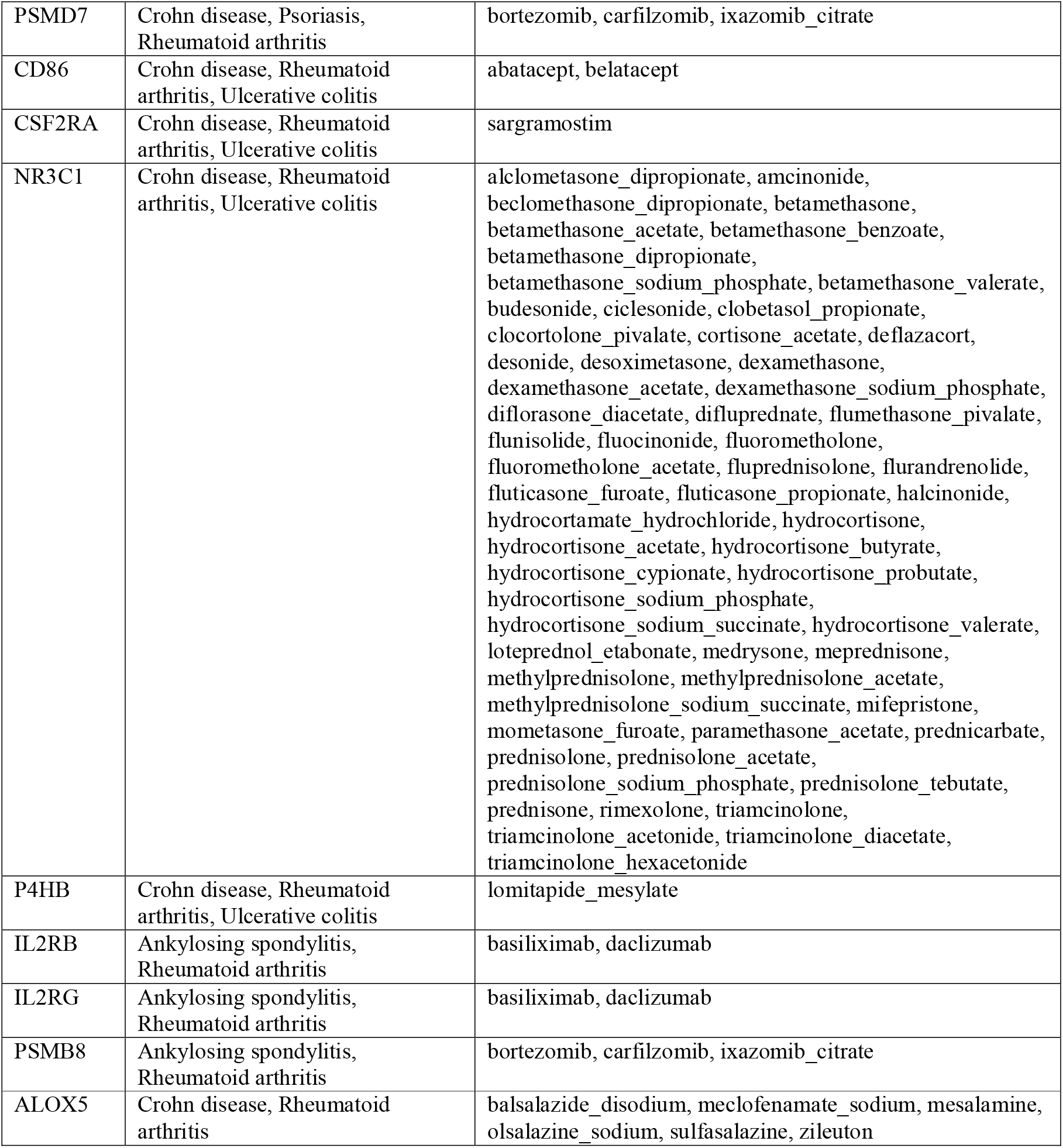
The top DAGs (from **Figure 5E-F**) that are drug targets along with their FDA approved drugs.

### 2.2. Disease-gene network from DisGeNet

The disease-gene network from DisGeNet [23] was downloaded from the DisGeNet database (www.disgenet.org/downloads). All HLA associated genes was removed from the network, this was done to ensure that bias towards myeloid cells and B cells are removed, since the HLA genes are largely expressed by these cells. The resulting network was further filtered to include only those genes that were present in the *immunome*.

### 2.3. IMID disease-gene network

To study and identify the DACs of the IMIDs, the DAGs of 12 IMIDs were extracted from the above DisGeNet. The IMID gene network for the 12 diseases comprised of 3579 DAGs. The 12 diseases that broadly represent the IMIDs in this study include: ankylosing spondylitis (CUI: C0038013), arthritis (CUI: C0003864), Crohn’s disease (CUI: C0010346), diabetes mellitus-non-insulin-dependent (CUI: C0011860), systemic lupus erythematosus (CUI: C0024141), multiple sclerosis (CUI: C0026769), psoriasis (CUI: C0033860), psoriatic arthritis (CUI: C0003872), rheumatoid arthritis (CUI: C0003873), Sjogren's syndrome (CUI: C1527336), systemic scleroderma (CUI: C0036421), and ulcerative colitis (CUI: C0009324). CUI, used in DisGeNet, is the concept unique identifier for the disease term as defined by unified medical language system [25]. The disease term arthritis (CUI: C0003864) comprises DAGs that pan over several arthropathies such as spondyloarthropathy, osteoarthritis, gout, allergic arthritis, etc., that fall under the broad arthritis MeSH term.

### 2.4. Identification of top DAC and DAG using machine learning

We used an unsupervised machine learning algorithm called non-negative matrix factorization (NMF) to map the disease-gene network to the *immunome*, and identify the top DACs and DAGs of the 12 IMIDs. The NMF algorithm clusters the input gene expression data into ‘k’ clusters, such that the DAGs of a cluster are expressed by the DACs of the same cluster, thus forming DAC-DAG pairs in each cluster [24]. We used the coefficients and weights identified by the NMF algorithm as the DAC and DAG scores respectively. The scores were scaled between 0 and 1, with 1 being the highest score. Those in the top 25 percentile of the scores were regarded as the top DACs and DAGs respectively. We calculated the Frobenius norm for each cluster to weigh and rank the clusters, the rank 1 cluster is referred to as the top cluster. The top cluster comprise the DAC-DAG pair that which maximally captures/represents the input gene expression matrix. Using the top DAC-DAG pairs of all clusters, we constructed the Disease-gene IMmune cell Expression (DIME) network for the 12 IMIDs (see Supplementary methods for details).

### 2.5. Common cell-gene network between diseases

To identify common cell-gene network between two diseases, we looked at their overlapping DAC-DAG pairs in their corresponding DIME networks. These overlapping DAC-DAG pairs are referred to as the common cell-gene network between the two diseases. Jaccard index (JI) was used to measure the overlap between the two diseases with Fisher’s exact test (FET) used to obtain confidence p-value for the given overlap.

### 2.6. Integrating drug-gene network

The drug-gene target network was curated from (1) DGIdb with the filter set to contain CHEMBL interactions pertaining to the drugs approved by the food and drug administration (FDA) of USA [26]; (2) all drug-gene of CLUE database [27] and; (3) all drug-gene of hPDI [28]. The genes that had drugs associated to them are labelled in the common cell-gene networks to highlight druggability (Figure 5C-E).

### 2.7. Statistical analysis

We performed 1000 jackknife simulations to assess the consistency of the results from the DIME (Supplementary methods, and Supplementary figure 1–3). Pearson correlation coefficient and p-value were computed to measure significance of the jackknife simulations in comparison to the original run (Supplementary figure 3).

## 3. RESULTS

### 3.1. Disease-gene network of the 12 IMIDs reveal several common DAGs

In this study, we analyzed different types of IMIDs that include inflammatory arthropathies, spondyloarthropathies, rheumatic diseases, systemic IMIDs, and inflammatory bowel diseases (IBDs). The 12 different IMIDs include: ankylosing spondylitis (DAGs:298), arthritis (DAGs:567), Crohn’s disease (DAGs:786), diabetes mellitus - non-insulin-dependent (DAGs:1415), systemic lupus erythematosus (DAGs:963), multiple sclerosis (DAGs:961), psoriasis (DAGs:689), psoriatic arthritis (DAGs:177), rheumatoid arthritis (DAGs:1612), Sjogren's syndrome (DAGs:229), systemic scleroderma (DAGs:494), and ulcerative colitis (DAGs:796) (Figure 1 A-B). In total, 3957 DAGs were linked to the 12 IMIDs. Among which, several genes were found to be linked to several IMIDs, for example, 74 DAGs were linked to only Crohn’s disease (CD) and to ulcerative colitis (UC), both IBDs (Figure 1A). Calculating the Jaccard index and Fisher’s exact test (FET) on all the overlapping DAGs between all IMIDs revealed that CD and UC had the highest significant overlap (Figure 1C). Interestingly, genes associated with CD had significant overlap (FET p-value ≤ 0.05) with all diseases except psoriatic arthritis and diabetes mellitus non-insulin dependent (T2D). Rheumatoid arthritis (RA) had significant overlap of DAGs with all IMIDs except T2D. T2D did not have significant overlap of DAGs with any of the IMIDs. Arthritis, psoriasis, CD, and RA had significant overlap of DAGs between each other. We found 12 DAGs that were associated with all the 12 IMIDs (Figure 1A, E). These DAGs were related to processes typically associated with inflammation such as: cytokine signaling (GO:0001817; GO:0019221), regulation of inflammatory response (GO:0050727), and regulation of interleukin-6 (GO:0032675; GO:0032635). We further explored the expression of these DAGs in the *immunome* and found the expression of TNFAIP3 to be the highest in CD8^+^ T-cells, ILC3 and CD4^+^ T-cells (Figure 1D, E). Likewise, IL1B was expressed by myeloid and progenitor cells; TNF was expressed by lymphoid and myeloid cells. Overall, certain myeloid cells and lymphoid cells, specifically expressed some of the 12 genes that were linked to all the 12 IMIDs. This intrigued us to identity the key immune cell types and genes that are important for the 12 IMIDs. Hence, we used the DIME on the 12 IMIDs to identify their top DACs and DAGs. Briefly, DIME uses the *immunome*, input disease-gene network and an unsupervised machine learning algorithm (NMF) to identify the clusters of top DACs and DAGs, see methods.

**Figure 1.**
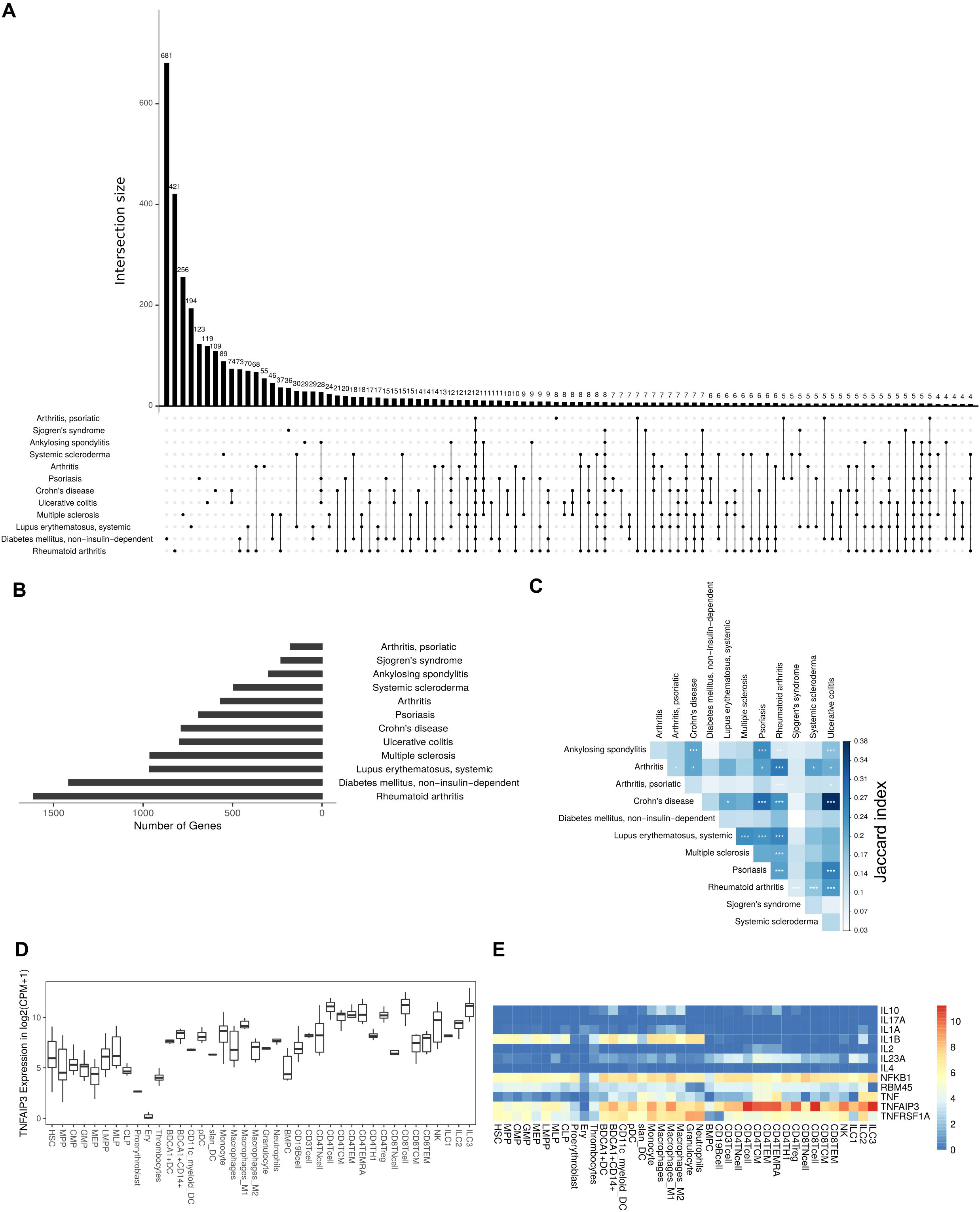
DAGs of IMIDs: **A**. UpSet plot showing the intersection of DAGs of all comparisons of IMIDs. Comparisons shown only for those disease that have at least 1 intersecting DAG between them. **B**. Barplot showing number of DAGs in each IMID. **C**. Heatmap depicting Jaccard index and Fisher exact test (FET) p-value calculated for each IMID comparison. Fisher exact test (FET) p-value denoted by * (*** ≤ 0.001, ** ≤ 0.01, and * ≤ 0.05). **D**. Gene expression of TNFAIP3. **E**. Heatmap depicting gene expression of the 12 genes common to all 12 IMIDs. Gene expression values are measured in log2(cpm+1), cpm denotes counts per million.

### 3.2. Top immune cells of inflammatory arthritis

Inflammatory arthritis is characterized by joint inflammation due to autoimmunity. Joint inflammation is the primary clinical feature as observed in ankylosing spondylitis (AS) and RA. However, in other inflammatory arthritis such as the psoriatic arthritis, inflammation is present in both the skin and joints. Interestingly, AS and psoriatic arthritis are both seronegative spondyloarthropathies (negative for rheumatoid factor and auto nuclear antibodies) that are characterized by enthesitis and also have a predominant HLA-B27 genotype [29,30]. We questioned if the inflammatory arthritis shared molecular mechanism, in addition to sharing clinical features. So, we performed DIME on the different types of inflammatory arthritis to identify the important DACs and DAGs, and compare the molecular mechanism shared between them. As a reference, we used the broader arthritis disease term that encompassed (including inflammatory arthritis) several different kinds of arthropathies, see methods for disease description.

The DIME analysis of ankylosing spondylitis revealed lymphoid cells such as NK cells, ILC3, CD4^+^ T-cells (Th1, Treg, TEMRA) as the top DACs in the top cluster (Figure 2A). The top DAGs of the top cluster were associated with pathways such as interleukin signaling, antigen presentation, regulation of RUNX3, and BCR signaling (Figure 2E). The role of RUNX3 in NK cells, CD4^+^ and CD8^+^ T-cells has been reported to be important in AS [31]. In the second cluster, the top DACs included myeloid cells and the top DAGs were associated with pathways such as interleukin (IL-4, IL-10, IL-13) signaling, MAPK3 activation and MyD88 (Figure 2A, E). Thus, the key DACs of AS were found to be diverse as reported in the literature, however the top DACs according to DIME were NK cells, ILC3, CD4^+^ T-cells (Th1, Treg, TEMRA) [32].

**Figure 2.**
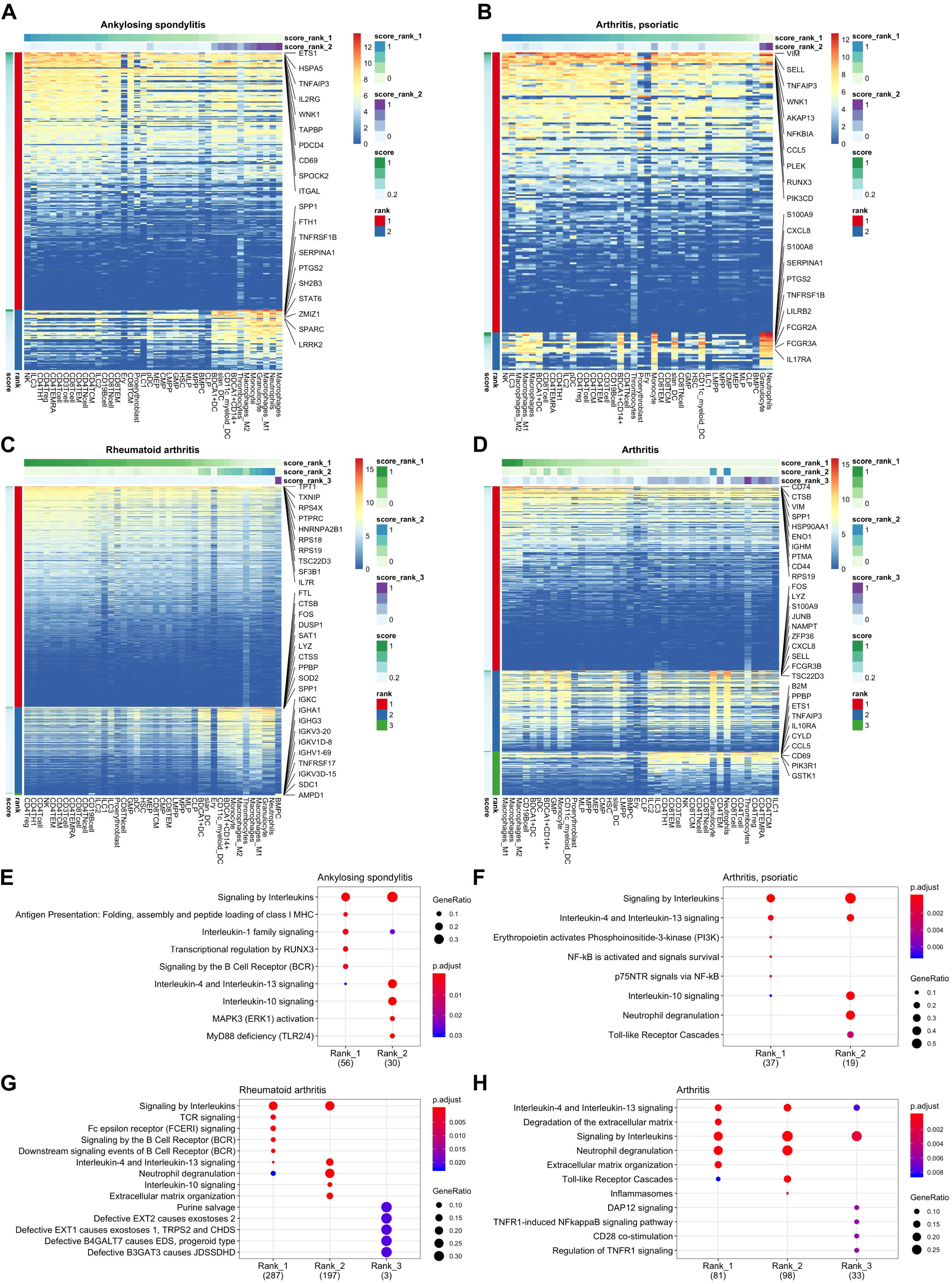
DIME analysis of inflammatory arthritis: DIME heatmaps and pathway enrichment analysis for the top 25 percentile DAG identified by DIME for ankylosing spondylitis (**A, E**), psoriatic arthritis (**B, F**), rheumatoid arthritis (**C, G**) and arthritis (**D, H**) respectively. The top 10 DAGs based on the DAG score are labelled in the DIME heatmap.

The DIME analysis of psoriatic arthritis revealed lymphoid cells such as NK cells, ILC3 and myeloid cells like the macrophages and BDCA1^+^ DC as the top DACs in the top cluster (Figure 2B). Likewise, T-cells, NK cells and antigen presenting cells have been reported to play a role in the pathology of psoriatic arthritis [33]. The top DAGs of the top cluster were associated with pathways such as interleukin (IL-4, IL-10, IL-13) signaling, activation of PI3K, and NF-KB. (Figure 2F). S100 calcium binding proteins like S100A8 and S100A9 are known to play a role in the regulation of inflammation in psoriatic arthritis [34]. In the second cluster, we found the top DAGs included the S100 calcium binding proteins, such as S100A9, and S100A8 that were highly expressed by the granulocytes, neutrophils, monocytes and dendritic cells (Figure 2B, F).

The major immune cells involved in RA are T-cells, B-cells, and APCs [35]. While activation of CD4^+^ Th1 and impairment of CD4^+^ Tregs have been reported to be important for rheumatoid arthritis [36], the DIME analysis of RA revealed several lymphoid cells such as CD4^+^ Tregs, CD4^+^ Th1, NK cells, etc., as the top DACs in the top cluster (Figure 2C). The top DAGs of the top cluster were associated with pathways such as interleukin, TCR, FCERI, and BCR signaling (Figure 2G). In the second cluster, the top DACs included myeloid cells and the top DAGs were associated with pathways such as interleukin (IL-10, IL-13) signaling, neutrophil degranulation, and ECM organization (Figure 2C, G). Evidently, activation, recruitment and apoptosis of neutrophils is altered in RA and under the chronic inflammatory conditions they release protease-rich granules [37].

The DIME analysis of the broader arthritis disease term, revealed macrophages as the top DAC in the top cluster (Figure 2D). Macrophages play a central role in arthropathies, where they release cytokines and activate several immune cells such as T-cells, monocytes, neutrophils, and synovial fibroblasts. In addition, they are also the most abundant cells at the site of inflammation [38]. The top DAGs of the top cluster were associated with pathways such as interleukin (IL-4, IL-13) signaling, extracellular matrix (ECM) related pathways, neutrophil degranulation and toll-like receptor (TLR) cascades (Figure 2H). In the second cluster, the top DACs comprise of neutrophils, granulocytes and the top DAGs were associated to pathways similar to the top cluster, and also included inflammasomes related pathways (Figure 2D, H).

### 3.3 Top immune cells of systemic IMIDs

We performed the DIME analysis on the systemic IMIDs such as systemic lupus erythematosus (SLE) and systemic scleroderma (SSc) (Figure 3). SLE and SSc are type I interferon-mediated systemic autoimmune diseases, that unlike RA, primarily affects not just the joints, but also the skin, kidney, heart, and other organs [39].In SLE, the continuous IFN production by pDC and neutrophils leads to activation of monocytes, T-cells, and B-cells [40]. The DIME analysis of SLE revealed the myeloid cells (granulocytes, macrophages, BDCA1^+^ CD14^+^, monocytes) as the top DACs in the top cluster (Figure 3A). The top DAGs in the top cluster were associated with pathways such as interleukin signaling (IL-4, IL-13), neutrophil degranulation, cell surface interactions at the vascular wall and the TLR cascades (Figure 3C). Incidentally, the neutrophils in SLE undergo spontaneous NETosis (a form of suicidal cell death) and this process is dependent on TLR signaling [40]. Additionally, T-cells in SLE are found to have altered cytokine production with higher levels of IL6, IL7, and IL10 secretions [40]. In the second cluster, we found the top DACs included CD4^+^ T-cells (TEMRA, TEM, TCM) and the top DACs were associated with pathways such as immunoregulatory interactions, Nef-associated factors (TNIP1, TNFAIP3), ZAP-70, VAV1 pathway (Figure 3A, C). Nef-associated factors (TNIP1, TNFAIP3) is known to play a role in activation of T-cell via TCR signaling in SLE [41].

**Figure 3.**
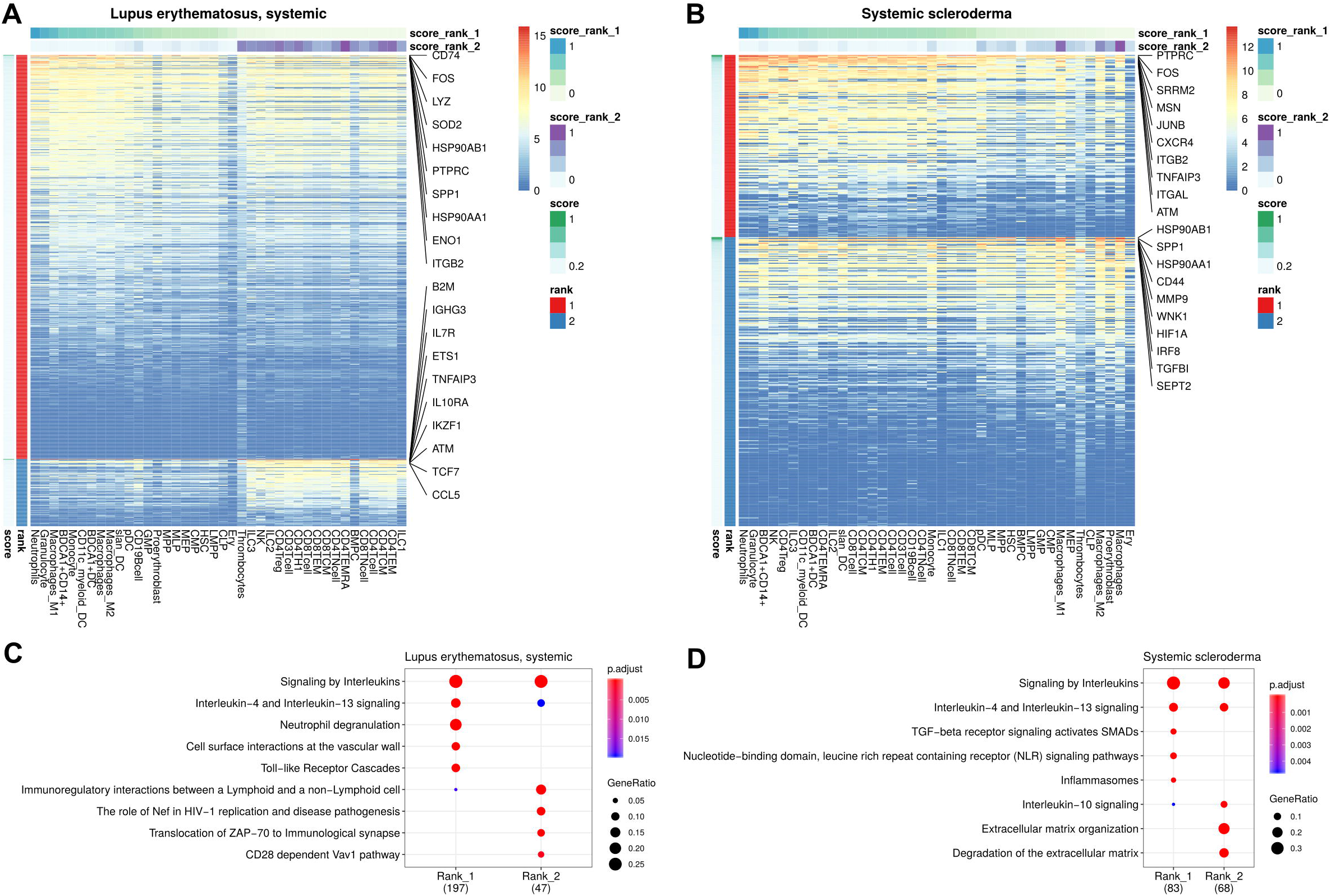
DIME analysis of systemic diseases: DIME heatmaps and pathway enrichment analysis for the top 25 percentile DAG identified by DIME for SLE (**A, C**), and SSc (**B, D**) respectively.

The DIME analysis of SSc revealed myeloid cells (neutrophils, granulocytes, BDCA1^+^ CD14^+^ cells) and lymphoid cells (NK cells and CD4^+^ Treg) as the top DACs in the top cluster (Figure 3B). The top DAGs in the top cluster were associated with pathways such as interleukin signaling (IL-4, IL-13), TGF beta signaling, NLR signaling, etc. (Figure 3D). In the second cluster, the top DACs included macrophages and the top DAGs were associated to pathways that included IL-10 signaling and degradation of ECM (Figure 3B, D). As described in the review by Caam et al., several studies have shown neutrophils, macrophages, NK cells, and Tregs to play a role in the profibrotic events in SSc by the production of profibrotic cytokines such as TGF beta, IL-4, IL-10, IL-13, etc., thus corroborating our findings [42].

### 3.4. Top immune cells in Inflammatory bowel diseases (IBDs)

We then looked at IMIDs that involve chronic inflammation of the digestive system, these are categorized as IBDs. The two major forms of IBDs are CD and UC. CD is known to be driven by CD4^+^ Th1 cells, with a dominant Th1 cytokine profile leading to pro-inflammatory effect [43]. The DIME analysis of CD revealed lymphoid cells (CD4^+^ Treg, ILC2, CD4^+^ TEMRA, CD4^+^ Th1) as the top DACs in the top cluster (Figure 4A). The top DAGs of the top cluster were associated with pathways such as interleukin (IL-4, IL-10, IL-13) signaling, TLR (TLR-5, TLR-10) signaling, MyD88 and neutrophil degranulation (Figure 4C). In the second cluster, the top DACs included granulocytes, neutrophils, monocytes, macrophages, etc., and the top DAGs were associated with pathways such as interleukin signaling, neutrophil degranulation and TLR cascades (Figure 4A, C).

**Figure 4.**
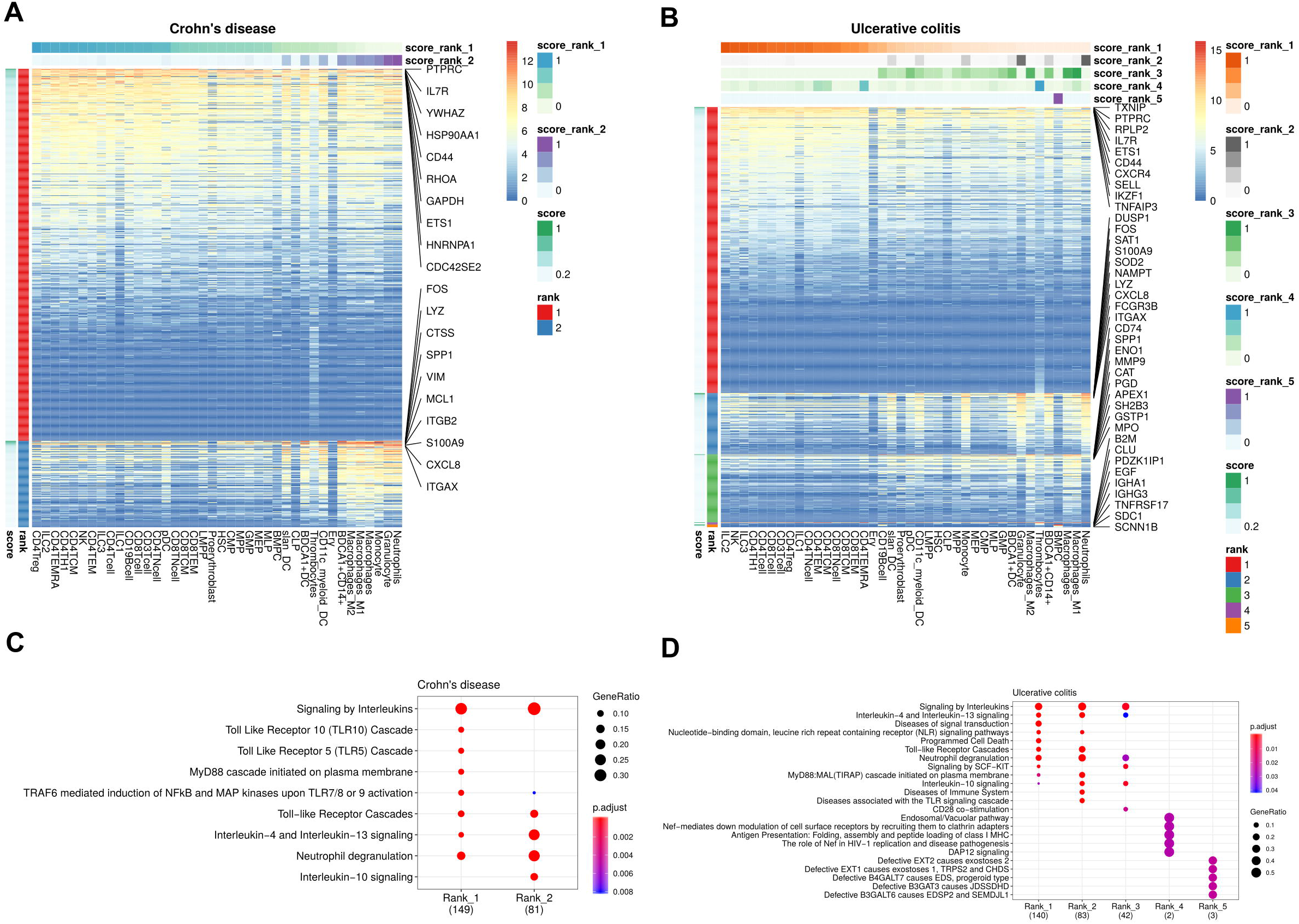
DIME analysis of IBDs: DIME heatmaps and pathway enrichment analysis for the top 25 percentile DAG identified by DIME for Crohn’s disease (**A, C**), and ulcerative colitis (**B, D**) respectively.

The T-cell profile of UC has been difficult to categorize due to discrepancies in its response among patients. However, there is evidence of Th2 cells, NK cells, macrophages, and neutrophils to be involved in the pathogenesis of UC [43]. The DIME analysis of UC revealed lymphoid cells (ILC2, NK, ILC3, CD4^+^ Th1, etc.) as the DACs in the top cluster (Figure 4B). The top DAGs of the top cluster were associated with pathways such as interleukin (IL-4, IL-13) signaling, TLR cascades, NLR signaling, neutrophil degranulation, etc. (Figure 4D). In the second cluster, the top DACs included granulocytes, BDCA1^+^ CD14^+^ cells, etc., and the top DAGs were associated with pathways such as interleukin signaling (IL-4, IL-10, IL-13), neutrophil degranulation and TLR cascades (Figure 4B, D).

### 3.5. Statistically significance of DIME results

To evaluate the consistency of results from DIME, we performed 1000 Jackknife simulations with random subsampling of DAC/DAG and re-identified the top DAC/DAG for all IMIDs (see Supplementary methods). The jackknife simulations revealed that the top DACs identified across all clusters in the simulations (Supplementary figure 1A) showed similar pattern when compared to top DACs identified in the original run (Supplementary figure 1C). For the top DACs of the top cluster, the pattern from the simulations (Supplementary figure 1B) were comparable to the DAC score of the original run (Supplementary figure 1D). We used Pearson correlation to compare the pattern observed between the simulations and the original run, see Supplementary methods. The Pearson correlation between the pattern observed in simulated run (Supplementary figure 1B), and the DAC scores of the original run for the top cluster revealed that the top DACs in top cluster were significantly correlated (p-value ≤ 0.05) for all the IMIDs except ulcerative colitis (Supplementary figure 3A). This shows that the top DACs of the top cluster identified by DIME are statistically significant for all IMIDs, except UC.

Likewise, we evaluated the consistency of the top DAGs. In all simulations, the top 10 DAGs of top cluster of the original run were present as the top DAG in any of the clusters of the simulated run (Supplementary figure 2). The presence of the top 10 DAGs of top cluster of the original run as the top DAG in the top cluster of the simulations was also found to be high. The Pearson correlation between the pattern observed in the simulated run and the DAG scores of the original run for the top cluster were found to be significantly correlated for all the IMIDs (Supplementary figure 3B, see Supplementary methods). This shows that the top DAGs of the top cluster identified by DIME are statistically significant for all IMIDs.

### 3.6. Why are the top DACs of UC insignificant?

In the case of UC, the top DACs was found to be statistically insignificant from our 1000 jackknife simulations, the top DAGs however, were significant (Supplementary figure 1-3). We found from 1000 simulations that the lymphoid cells identified by the original run (Figure 4B) were indeed present in the simulations, and in addition, the myeloid cells were also part of the top DACs of the top cluster in the simulations (Supplementary figure 1B). Furthermore, we found that the top DAGs of the top cluster included genes associated to neutrophil degranulation pathways and other myeloid cell related pathways (Figure 4B, D), thus, owing to the non-convergence of the NMF algorithm in accurately predicting the top DACs of the top cluster in UC. The top DACs of the top cluster of UC was found to be ambiguous as has been reported in the literature [43]. From our simulations, we propose the inclusion of the myeloid cells in the top DACs of the top cluster in addition to the lymphoid cells previously identified (Figure 4B).

### 3.7. Common mechanisms in IMIDs

The DIME analysis revealed that several top DAGs along with their corresponding DACs were present in many IMIDs. For example, in many IMIDs, the gene FOS was present as top DAG in the cluster typically containing myeloid cells (granulocytes, neutrophils and dendritic cells) as the top DACs. We found several genes like FOS, that were present as the top DAG in the same top DAC cluster between different pairs of diseases. We refer to these top DACs and DAGs that are present between the two diseases as the common cell-gene network, represented schematically in Figure 5A, see methods. Using, the common cell-gene network, we suggest that these diseases may have similar mechanism of action. Such common mechanisms can be exploited to gain mechanistic insights between diseases and to identify drug repurposing targets. Hence, we integrated the drug-gene networks (see methods) to identify and reinforce drug repurposing targets based on the common mechanisms (cell-gene networks) identified from the DIME analysis, Figure 5A.

**Figure 5.**
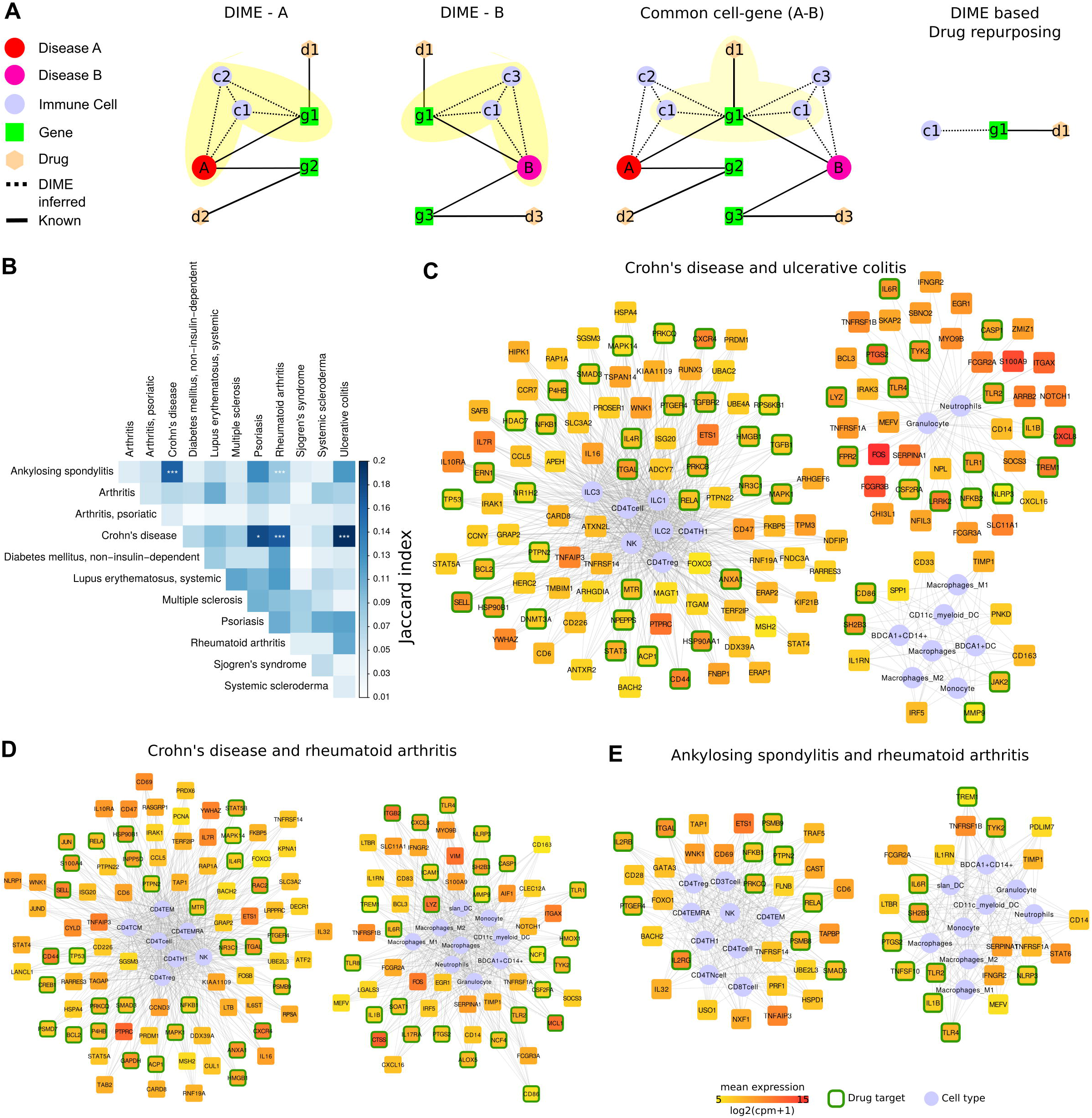
Common mechanisms between IMIDs. **A**. Steps involved in DIME based drug repurposing using the common cell-gene network. **B**. Jaccard index and FET calculated for the common cell-gene between two diseases for all disease comparisons. Fisher exact test (FET) p-value denoted by * (*** ≤ 0.001, ** ≤ 0.01, and * ≤ 0.05). Common cell-gene network of **C**. Crohn’s disease and ulcerative colitis. **D**. Crohn’s disease and rheumatoid arthritis. **E**. ankylosing spondylitis and rheumatoid arthritis. The cells are shown in blue. Color of the DAG is based on the mean gene expression of the DAG in the corresponding DACs. DAGs that are drug targets have a green border.

To identify the common mechanisms across the 12 IMIDs, we identified the common cell-gene networks between all disease comparisons (Figure 5B). Jaccard index and FET was used to measure the extent and significance of the overlap in the common cell-gene networks between the different pairs of diseases, respectively, see methods. In comparison to the analysis that looked at all DAGs, which showed several diseases to be statistically significant in the overlap (Figure 1C), the common cell-gene network overlap was restricted to fewer diseases (Figure 5B).

The comparative analysis revealed that CD had statistically significant common cell-gene networks with several diseases such as AS, psoriasis, RA, and UC (Figure 5B). Among which, the common cell-gene network of CD and UC had the highest Jaccard index among all the IMIDs, both being IBDs with aggressive T-cell response [43]. The common cell-gene network of CD and UC revealed that the top DACs included the lymphoid cells such as CD4^+^ T-cell, CD4^+^ Th1, CD4^+^ Treg, ILC1, ILC2, ILC3 and NK cells in one cluster (Figure 5C). CD4^+^ Th1 and NK cells are known to be implicated in both CD and UC [43]. The top DAGs such as CXCR4, IL10RA, IL7R, ETS1, TNFAIP3, PTPRC, SELL, etc., were highly expressed by cells of the lymphoid cluster. These DAGs were enriched in pathways associated with interleukin signaling (IL-4 and IL-13), NLR signaling, etc (Supplementary figure 4A). The other clusters comprised of myeloid cells such as the granulocytes, dendritic cells, monocytes and macrophages, among which dendritic cells have been crucial for regulating the T-cell responses in IBDs. The top DAGs such as IL6R, CXCL8, ITGAX, S100A9, FOS, etc., were highly expressed by the cells of the myeloid cluster. These DAGs were enriched in pathways associated with interleukin signaling (IL-10), TLR signaling, ECM degradation, etc (Supplementary figure 4A).

We next explored the common cell-gene network of the two distinct IMIDs that belonged to different pathophysiology, namely CD and RA. The common cell-gene network of CD and RA revealed that the top DACs comprised of the lymphoid cells that included all CD4^+^ T-cells and NK cells in one cluster (Figure 5D). The top DAGs such as CD69, PTPRC, CXCR4, etc., were highly expressed by the cells of this cluster. These DAGs were enriched for pathways associated with interleukin, TLR, MyD88 signaling, etc. (Supplementary figure 4B). The other clusters comprised of myeloid cells such as the granulocytes, dendritic cells, monocytes and macrophages. The top DAGs such as CTSS, ITGB2, ITGAX, MCL1, FOS, etc., were highly expressed by the cells of this cluster. These DAGs were enriched for pathways associated with interleukin (IL-4, IL-13) signaling, neutrophil degranulation, etc. (Supplementary figure 4B).

In addition to the common cell-gene networks of CD, we also found statistically significant common cell-gene network between the two inflammatory arthropathies that has joint pain as the primary feature, namely AS and RA. The common cell-gene network of AS and RA revealed that the top DACs comprised of the lymphoid cells that included all the T-cells, and NK cells in one cluster (Figure 5E). The top DAGs such as ITGAL, ETS1, IL2RG, TNFAIP3, etc., were highly expressed by the cells of this cluster. These DAGs were enriched for pathways associated with interleukin (IL-1) signaling, FCERI mediated NF−kB activation, TCR signaling, etc. The other clusters comprised of myeloid cells such as the granulocytes, dendritic cells, monocytes and macrophages. The top DAGs such as TNFRSF1B, STAT6, TYK2, were highly expressed by the cells of this cluster. These DAGs were enriched for pathways associated with interleukin (IL-4, IL-10, IL-13) signaling (Supplementary figure 4C).

Thus, using the common cell-gene networks we were able to uncover the common mechanisms in accordance to the top DACs and DAGs (Figure 5). This revealed several pathways that are common between the different IMIDs (Supplementary figure 4). Our next question was to see if these common mechanisms comprised of any drug targets (genes that are druggable or have drugs that target them). The idea was to identify targets for drug repurposing based on the drug targets in the common cell-gene networks, this novel method of computational drug repurposing is a combination of target-based and mechanism-based drug repurposing strategies [44]. To perform this, we then used the common cell-gene networks identified here and the drug-gene networks from literature to explore the common DAGs that were also drug targets (Figure 5C-E, DAGs highlighted by green border), see methods. We found several DAGs such as IL1B, IL6R, ITGAL, PTGS2, TYK2, NFKB1, NLRP3, PRKCQ, PTGER4, PTPN2, RELA, SH2B3, SMAD3, TLR2, TLR4, and TREM1, that were drug targets and present in all the common cell-gene networks shown in Figure 5C-E. Interestingly, ITGAL was found to be the only DAG that was a drug target and present as the top DAG of the top cluster (lymphoid cell cluster) in the DIME networks of CD, UC, AS and RA. Using the drugs associated to these drug targets specifically for these diseases (CD, UC, AS, and RA) in therapy would require extensive experimental validation and clinical trials. Therefore, we explored (in the next section) the possibility of using some of these drug targets for repurposing based on existing studies. Thus, reinforcing and strengthening these targets and also the validity of our approach in identifying them.

### 3.8. Common cell-gene networks from DIME reveals drug targets for repurposing

To explore and validate the drug targets for repurposing, we focused on the top DAGs of the statistically significant (FET p-value Ill 0.05, Figure 5B) common cell-gene networks of all IMIDs. To identify drug targets that were targets of FDA approved drugs, we used the drug-gene network of CHEMBL, see methods. We found several drug targets (Table 1) such as IL1B, IL6R, ITGAL, and TYK2 to be present in all the statistically significant common cell-gene networks. Anti-IL1 therapy is used for psoriasis and RA [45–47]. Preliminary studies indicate that anti-IL1 therapy has shown promising clinical response for treating AS, CD, and UC [48,49]. Anti-IL6 therapy (tocilizumab) shows positive clinical response in small group of patients in AS [50], CD [51] and in RA [52]. However, anti-IL6 therapy was found to have side effects in smaller studies on psoriasis and UC [53,54]. Integrin based therapies (such as natalizumab and vedolizumab that targets ITGB2) are already in use for CD [55]. Exploring other integrin based therapies (such as Lifitegrast that targets ITGAL and also ITGB2) for CD may be beneficial since both ITGAL and ITGB2 are top DAGs and are also implicated in CD [56,57]. Lifitegrast is a promising drug repurposing candidate for CD and also perhaps for UC, AS, and RA, since its target gene ITGAL, was the only top DAG of the top cluster (lymphoid cell cluster) that was also a drug target in the DIME networks of these diseases (Figure 2, 4, 5C-E). Thereby, targeting the same mechanism implicated in these diseases.

Tofacitinib, a TYK2 and JAK2 inhibitor developed for RA is now making way to treatment options in other diseases such as AS, CD, UC and psoriasis [58–61]. Corticosteroids (drug target: NR3C1) and the aminosalicylates (drug target: PTGS2 and ALOX5) are current line of drugs used in treatment of several IMIDs [62]. Plerixafor (drug target: CXCR4) is a drug currently used in cancer (lymphoma and multiple myeloma), after stem cell transplantation to initiate migration of stem cells in the bloodstream [63]. This drug is now in clinical trials (NCT01413100) to be evaluated for use after autologous transplant in patients with SSc. Such trials may potentially be extended to other IMIDs like psoriasis, CD, RA, and UC, that are driven by CXCR4 mediated dysregulation of immune system.

## 4. Discussion

Despite decades of experimental data, the knowledge on key cell types that are involved in pathogenesis of the disease still remains limited. To address this gap, we used the *immunome* comprising 40 immune cells, the disease-gene network and computational methods to identify the important DACs and DAGs of the disease. The integration of these parts resulted in the novel mechanisms being captured by our method, using which we built a tool called the DIME. Here, we highlight the important DACs, DAGs, and common mechanisms captured using DIME for 12 phenotypically different IMIDs. Using DIME, the top DACs were found to be CD4^+^ Treg, CD4^+^ Th1, and NK in the inflammatory arthritis (AS, PsA, and RA); neutrophils, granulocytes and BDCA1^+^CD14^+^ cells in SLE and SSc; ILC2, NK, CD4^+^ Th1, and CD4^+^ Treg in the IBDs.

Lymphoid cells such as CD4^+^ Th1, CD4^+^ Treg and NK cells were found to be the key players in inflammatory arthritis (AS, PsA and RA) and IBD (CD and UC). These diseases have been reported to have an intricate cross play of the above lymphoid cells, where the NK cells influence the differentiation of CD4^+^ Th cells into CD4^+^ Th1 and CD4^+^ Tregs; CD4^+^ Th1 plays a key role in initiation of inflammation by cytokine production; the CD4^+^ Tregs are crucial for immune response modulation [64]. Interestingly, the top DAGs of these diseases show pathways associated to signaling of IL-4 and IL-13 that are crucial in this cross play, thus corroborating the results from DIME.

Although, our analysis excluded HLA genes to avoid myeloid and B cell bias, the IMIDs associated with the HLA-B27, such as psoriasis, AS, and IBDs were found to have statistically significant common-cell gene networks. However, PsA (also associated with HLA-B27) is not included here, since it did not have statistically significant common-cell gene network with any of the IMIDs (Figure 5B). Additionally, AS and RA, the two inflammatory arthritis with joint inflammation as the primary feature, also had statistically significant common-cell gene network. Thus, showing that the diseases with these shared clinical features also had common mechanisms as identified by DIME. The common mechanisms from these networks revealed several lymphoid and myeloid cells, and their expressing DAGs. The lymphoid cells such as CD4^+^ Th1, CD4^+^ Treg, and NK was predominant in all the statistically significant common-cell gene networks, showing that these diseases were indeed driven largely by the aggressive T-cell response [31–33,43]. Pathways such as interleukin (IL-4 and IL-13), TLR, TCR signaling, etc., was found to be enriched in the top DAGs of the common cell-gene networks of these IMIDs. Thus, the common cell-gene network revealed several common mechanisms between the diseases in accordance to the top DACs, DAGs, and their associated pathways.

We used the information of the common mechanism from the common cell-gene network and the drug-gene networks to propose potential drug targets for repurposing. This novel computational drug repurposing strategy, a combination of target-based (literature drug-gene network) and mechanism-based (inferred from DIME) revealed several potential drug targets such as IL1B, IL6R, ITGAL, PTGS2, TYK2, NFKB1, NLRP3, PRKCQ, PTGER4, PTPN2, RELA, SH2B3, SMAD3, TLR2, TLR4, and TREM1. Further, we used these mechanism-based drug targets from DIME and the FDA approved drug-gene network to propose several drug targets and their drugs that could expedite the drug repurposing process (Table 1). Thus, we were able to capture drugs targets and their drugs that are currently being targeted or being explored for use in therapy for the IMIDs. We also found a few novel targets such as the drug lifitegrast (used for dry eyes) for CD, UC, AS and RA as an alternative to other integrin-based therapies already in use for CD. Lifitegrast is particularly interesting because it targets ITGAL, which was found to be important in the lymphoid cell cluster of CD, UC, AS and RA. Thus, effectively targeting the same mechanism. Perhaps the effect of lifitegrast on down-regulating lymphoid cell mediated inflammation [65] could be used in these diseases. Although, Lifitegrast is currently available as eye drops and used to treat only eye complications, different formulations of this drug can be explored to treat CD, UC, AS and RA. So far, to our knowledge, the use of drug lifitegrast in the axis of ITGAL, for the treatment of CD, UC, AS and RA has not been explored. Thus, using DIME, we were able to propose a novel drug repurposing strategy from the analysis of the 12 IMIDs.

## 5. Conclusions

Thus, DIME was helpful in identifying: 1. top DACs, DAGs of the IMIDs, 2. Common mechanisms between the IMIDs, and 3. drug targets for repurposing. To enable DIME analysis for other diseases from the DisGeNet, the GWAS network and also for user defined set of genes, we built the DIME tool as a user-friendly shinyapp. We believe that this tool will aid scientist to increase the understanding of disease pathology and facilitate drug development by better determining drug targets, thereby mitigating risk of failure in late clinical development.

## Data availability

All data presented here are available from the corresponding author upon reasonable request.

## Code availability

DIME tool is available on https://bitbucket.org/systemsimmunology/dime.

## Author contributions

AD and AP were involved in the conception of the study. AD was involved in the data curation, visualization and R shiny package development. AD and AP were involved in the data analysis and interpretation. AD and AP drafted the manuscript. TR helped in writing and revising the manuscript, and discussions about clinical perspective. All the authors revised the manuscript critically for important intellectual content and approved the submitted version.

## Acknowledgements

The authors thank Ajinkya Kadu for sharing his expertise in linear algebra and multivariate analysis. AP would like to acknowledge the Netherlands Organization for Scientific Research (NWO; Grant number 016.Veni.178.027) for financial support.

---

**Supplementary Table 1**: GEO datasets and samples used to construct the immunome.

