## Supplementary for "Integration of immunome with disease-gene network reveals common cellular mechanisms between IMIDs and drug repurposing strategies"

### Supplementary methods

#### Transcriptome data - *Immunome*

All the selected datasets (Supplementary Table 1) were downloaded as FASTQ files using the fastq-dump tool from sratoolkit. The "split-files" option was given if the library type was paired end sequencing. FASTQ files were then aligned to reference genome (GRCH.Hg38.79) using STAR aligner [1]. The result is a SAM file which was then converted into a sorted BAM file using the samtools program [2]. These were then used to calculate the count of aligned reads using the HTSeq program [3] with the mode option "intersection non-empty". HTSeq was run for all possible stranded mode options, the count file with the maximum counts was chosen as the respective count file for the sample.

The data was then filtered by removing all genes that had less than 20 read counts in 95 percent of the samples using R programming. The filtered data was then lane normalized using the "betweenLaneNormalization" function from the RUVSeq package [4]. The RUVr method from RUVSeq was used to identify residual factors contributing to the batch effect. The resulting filtered, batch corrected and normalized data had expression for 34,906 genes that was void of any observable batch effect. We calculated counts per million (cpm) for the filtered genes and used log2(cpm+1) as the gene expression measure. We then used the median gene expression for each cell type for the rest of the analysis. This processed, batch corrected, normalized and median representative data of 40 immune cells is referred here as the *immunome.*

#### Mapping disease-gene network to *immunome* data

For a given disease $D$ and its DAG, we first extracted the corresponding *immunome* expression matrix ($X_D$). $X_D$ comprised the gene expression of the DAG across the 40 cell types. $X_D$ was used as input matrix for the NMF algorithm.

#### Using NMF to cluster $X_D$ into $k$ classes

We used the NMF package[5] in R and applied Brunet's NMF algorithm [6] on $X_D$ to factor it into two matrices namely $W_D$ and $H_D$ such that.

$$X_D \approx W_D H_D \text{ (Equation 1),}$$

$$W_D H_D = \begin{bmatrix} | & | & | & | \\ w_D^1 & w_D^2 & \dots & w_D^k \\ | & | & | & | \end{bmatrix} \begin{bmatrix} - & h_D^1 & - \\ - & h_D^2 & - \\ - & \vdots & - \\ - & h_D^k & - \end{bmatrix} \text{(Equation 2)},$$

$$W_D H_D = \sum_{i=1}^{k} w_D^i\, h_D^i;\ i \in \{1, \dots, k\} \text{ (Equation 3)}$$

where, $W_D$ and $H_D$ are the basis and coefficient matrices computed by NMF. $k$ is the number of classes/clusters that splits the data, such that it satisfies the above NMF equations. The $W_D$ matrix comprises of the weights of the DAGs across the $k$ clusters (in each column) and the $H_D$ matrix comprises of the weights of the cells in the corresponding $k$ clusters (in each row). We used Brunet's method to identify the ideal $k$ value using the cophenetic correlation coefficient [6].

#### Identifying the top DAG and DAC from $W_D$ and $H_D$

The NMF algorithm clusters the data into $k$ clusters (as shown in Equation 2 and 3) such that, in each cluster '$i$', where $i \in (1, \dots, k)$, the genes that have high values in $w_D^i$ are constitutively expressed by the cells that have high values in $h_D^i$. Where, $w_D^i$ is the $i^{th}$ column of $W_D$ and $h_D^i$ is the $i^{th}$ row of $H_D$. We used the scaled (between 0 and 1) values of $h_D^i$ and $w_D^i$ as the DAC and DAG scores respectively. For each cluster $i$, we chose the DACs and DAGs that were in the top 25$^{th}$ percentile range of their DAC and DAG scores respectively. These filtered DACs and DAGs are regarded as the top DACs and DAGs respectively. The top DACs and DAGs were extracted for all clusters of $i \in (1, \dots, k)$. The DIME network was constructed using the top DAC-DAG pairs from all clusters.

#### Identifying the top cluster

We then identified the largest weighted cluster (referred to as the top cluster) among the $k$ clusters identified by the NMF. That is, the subset of DACs and DAGs of $X_D$ that can capture most of its expression pattern. We did this by calculating the Frobenius norm of each $w_D^i h_D^i$ for all values of $i \in (1, \dots, k)$ from Equation 3. We then identified the top cluster for which $||w_D^i h_D^i||_F$ is the maximum. This can be represented as:

$$top\ cluster = \operatorname{argmax}\left(||w_D^i h_D^i||_F\right);\ i \in \{1, \dots, k\} \text{ (Equation 4)}$$

where, the top cluster represents that which maximally captures/represents the expression matrix $X_D$. Thus, the top cluster is the rank 1 cluster of DIME. Subsequent ranks are the next highest weighted clusters.

**Evaluating the consistency of top DACs and DAGs identified by DIME**

To check the consistency of the results from DIME, we performed 1000 jackknife simulations for each of the 12 IMIDs. For each simulation of each disease, we ran DIME with 70% random subsampling of either the DACs or the DAGs. And in each simulation, we identified the top cluster, and the top DACs when DAGs were subsampled and vice versa. We compared the consistency of the top DACs (Supplementary figure 1), and the top 10 DAGs (Supplementary figure 2) identified by the original DIME run (100% of sample) against the 1000 simulations. We computed the Pearson correlation coefficient between the DAC/DAG score of the top cluster of the original run to the number of times the DAC/DAG was found as the top DAC/DAG in the top cluster of the 1000 simulations. We used the p-value from the Pearson correlation test to state significance of correlation and thus statistical significance of the top DAC/DAG of the top cluster.

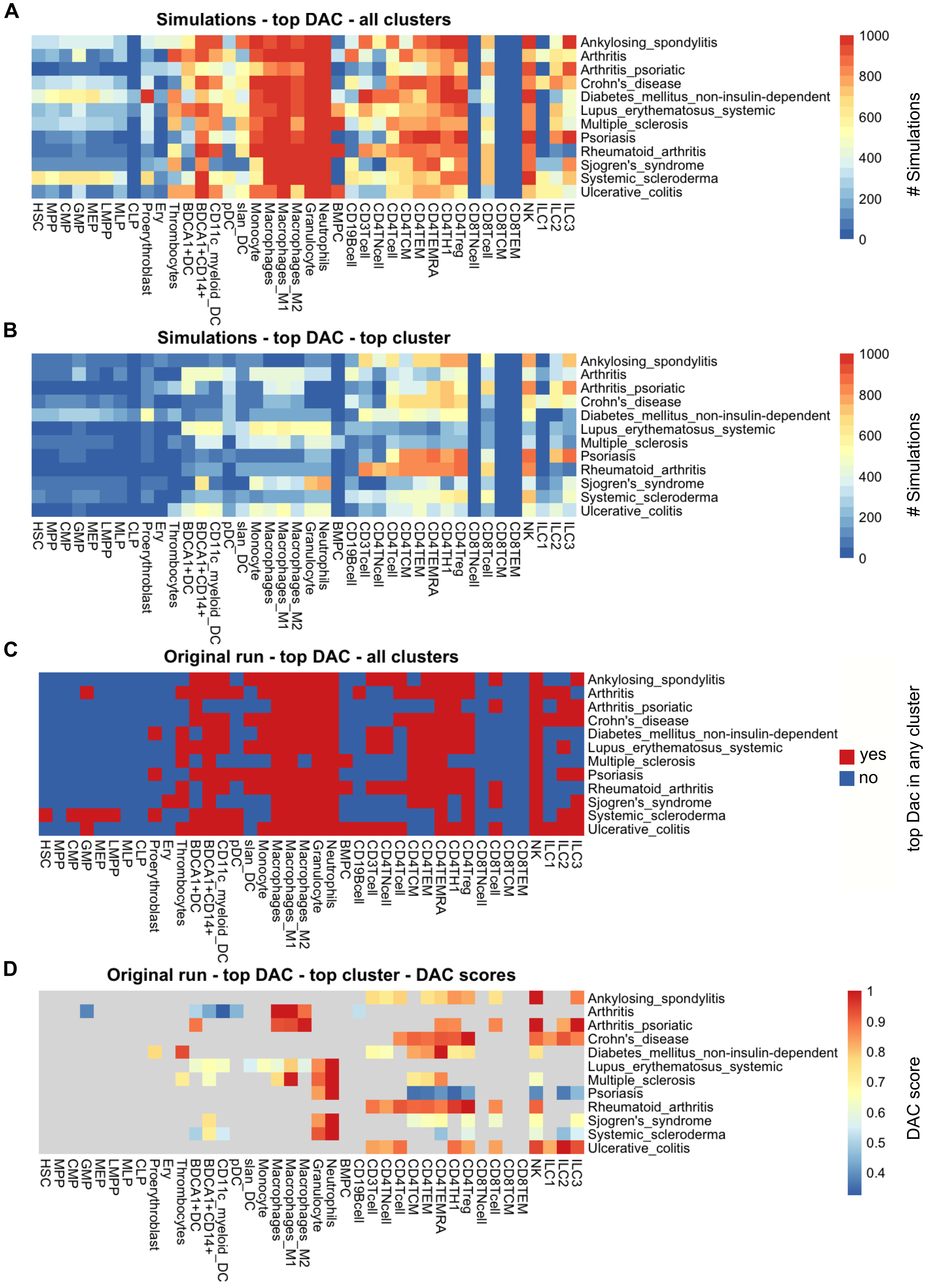


**Supplementary figure 1**: Consistency of top DACs: Heatmap of **A**. top DACs from all clusters **B**. top DACs from top cluster, identified using 1000 jackknife simulations with 70% random subsampling of DAGs. **C**. top DACs from all clusters of the original run, red represents if the cell type was a top DAC in any cluster. **D**. DAC score of the top DAC from the top cluster.

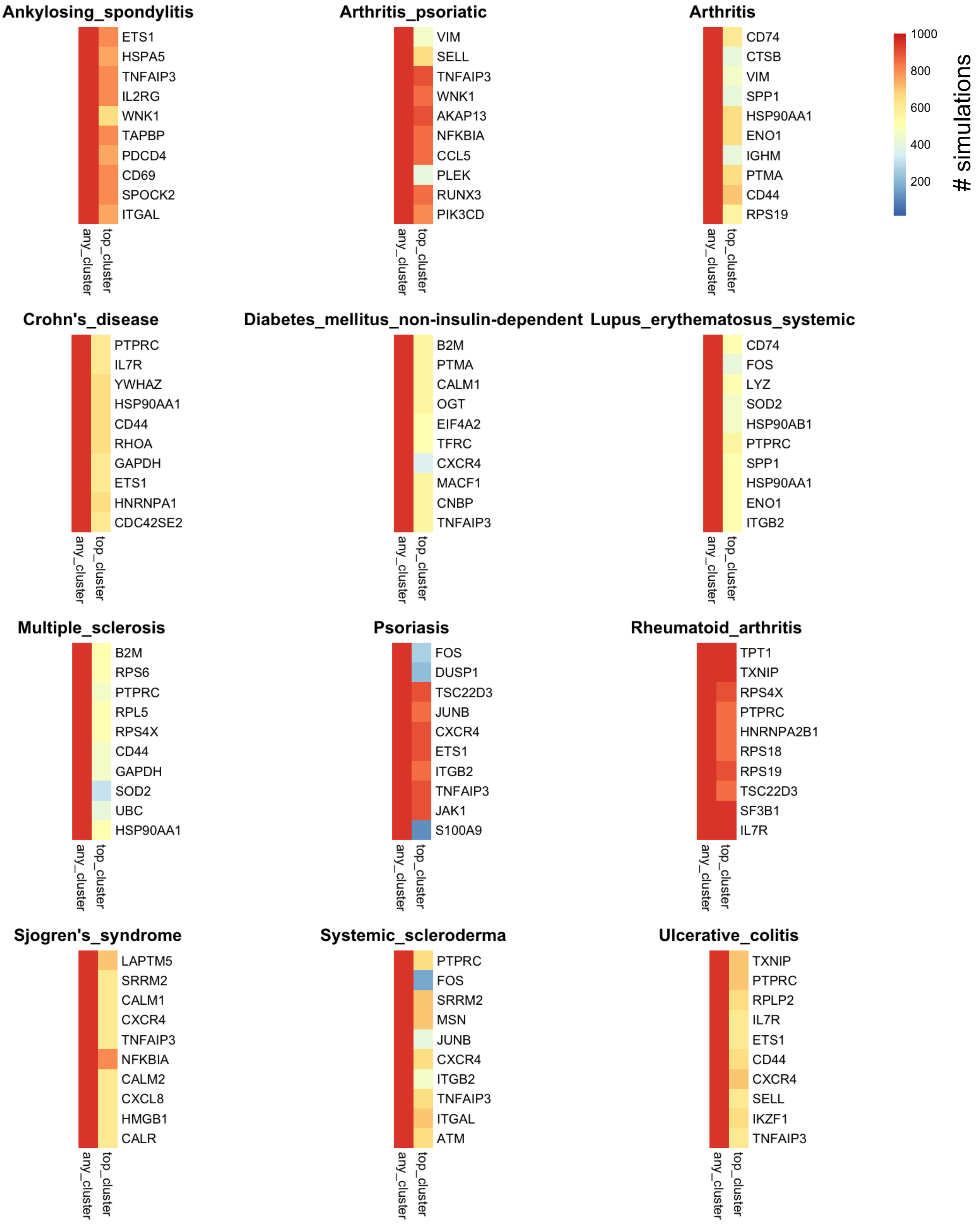


**Supplementary figure 2**: Consistency of top DAGs: Heatmap showing consitency of top 10 DAGs of the original run in the 1000 jackknife simulations with 70% random subsampling of cell types. Heatmap represents the numer of simulations in which, the top DAG was found in any cluster (column 1); was found in the top cluster (column 2).

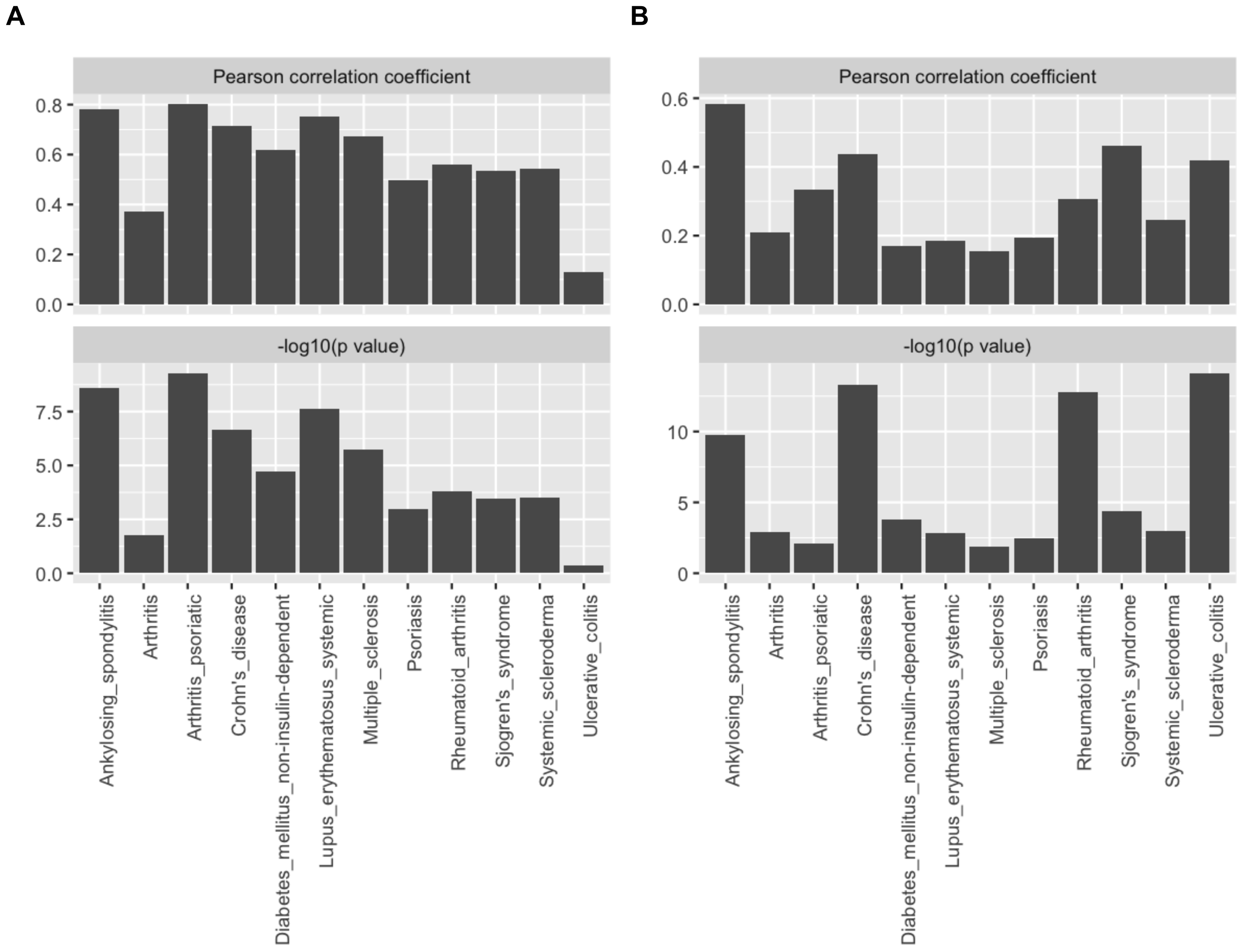


**Supplementary figure 3**: Pearson correlation coefficient and it's p-value calculated between the 1000 jacknife simulations and the **A**. DAC score, **B**. DAG score of the top cluster of the original run. p-value represented as -log10(p-value). -log10(0.05) = 1.301. The p-values of all the results were statisticall significant (-log10(0.05) ≥ 1.301) except the DAC simulations of ulcerative colitis.

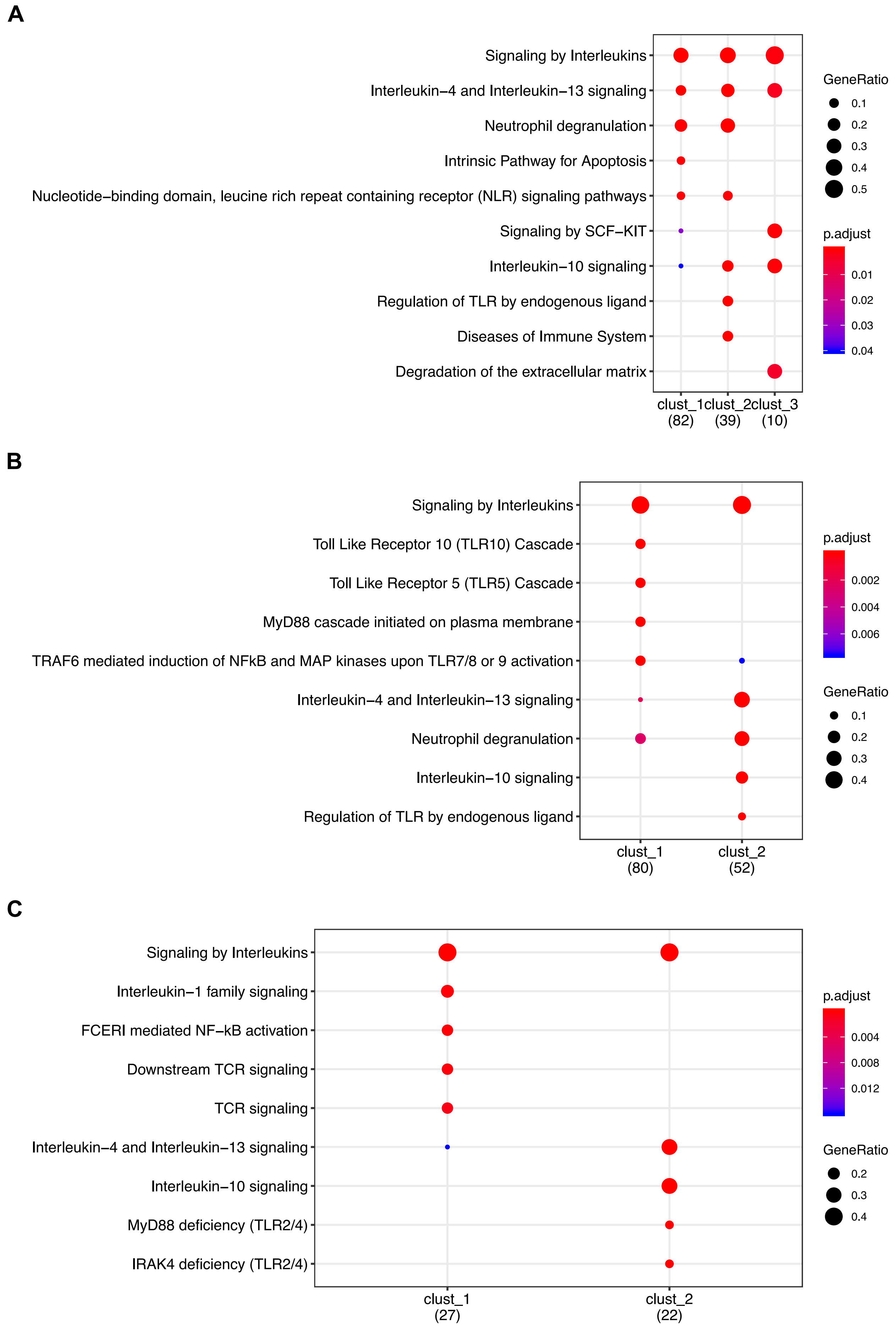


**Supplementary figure 4**: Pathway enrichment analysis of top DAGs of the common cell-gene networks between **A.** CD and UC; **B**. CD and RA; **C**. AS and RA.
